# Locomotor performance and CNS responses to hypoxia in a cyclic nucleotide-gated channel mutant of adult *Drosophila*

**DOI:** 10.1101/2021.02.11.430836

**Authors:** Shuang Qiu, Chengfeng Xiao, R Meldrum Robertson

## Abstract

*Drosophila* provides an excellent opportunity to explore the genetic basis for behavioral and CNS responses to hypoxia. Cyclic guanosine monophosphate (cGMP) modulates the speed of recovery from anoxia in adults and mediates hypoxia-related behaviors in larvae. Cyclic nucleotide-gated channels (CNG) and cGMP-activated protein kinase (PKG) are two cGMP downstream targets. PKG is involved in behavioral tolerance to hypoxia and anoxia in adults, however little is known about CNG channels. We used a CNGL mutant with reduced *CNGL* transcripts to investigate the contribution of CNGL to the hypoxia response. In control flies (w1118), hypoxia immediately reduced path length per minute in a locomotor assay. Flies took 30-40 mins in air to recover from 15 mins hypoxia. CNGL mutants had reduced locomotion under normoxia and impaired recovery from hypoxia, similar to the effects of pan-neural *CNGL* knockdown. In the CNGL mutants hypoxia caused an acute increase in path length per minute followed by a gradual increase during hypoxia. Basal levels of CNS extracellular K^+^ concentrations were reduced in the mutants. In response to hypoxia, the mutants had an increased extracellular K^+^ concentration change, reduced time to reach the K^+^ concentration peak, and delayed recovery time. Genetic manipulation to increase cGMP in the CNGL mutants eliminated the impairment of recovery from hypoxia and partially compensated for the effects of hypoxia on CNS K^+^. Although the neural mechanisms have yet to be determined, CNGL channels and cGMP signaling are involved in the hypoxia response of adult *Drosophila*.

## 1. Introduction

Insects have evolved remarkable adaptive mechanisms to tolerate hypoxia or anoxia. The median lethal time (LT50) of *Chironomus plumosis* larvae exposed to anoxia can be as long as ~200 days (48). The larvae of *Cicindela togata* survive anoxia at 25 °C with an LT50 of more than three days (24, 25). *Locusta migratoria* are able to survive exposure to 100% nitrogen for up to six hours at room temperature (70). Adults of *Drosophila melanogaster* can survive several hours in anoxia (5, 6, 8, 23). At extremely low oxygen tension (0.1 kPa), flies reduce metabolism around 10-fold relative to normoxia (64). The exceptional survival and physiological responses to hypoxia or anoxia in *Drosophila* provide a good opportunity to explore its underlying molecular and genetic basis (7).

The ability to sense oxygen levels is critical for eliciting long-term change and short-term behavioral responses to hypoxia and anoxia (38). Prolonged or chronic responses to hypoxia, such as hypoxia-induced gene expression, are primarily mediated by the conformational stabilization of hypoxia-inducible factors (HIFs) (26, 29, 57). However, rapid behavioral and electrophysiological responses to hypoxia often involve the activation of oxygen sensors and oxygen-operated ion channels (29, 33).

Ion channels can be regulated by reduced O_2_ levels (35) and the first identified ion channel responsible for O_2_ sensing was a voltage-dependent K^+^ channel (37). It is found in Type I chemoreceptor cells in the mammalian carotid body and hypoxia (*P*O2 dropping from 150 to 10 mm Hg) reduces the K^+^ current through the channel by 25-50 %. Another O_2_-sensing channel is the O_2_-sensitive ATP-inhibitable K^+^ channel, which is found in neocortical and substantia nigra neurons in the rat CNS (27, 28). Hypoxia activates this channel and causes it to open and close with increased frequency. Moreover, in rat hippocampal CA1 neurons, the voltage-dependent, fast-inactivating Na^+^ inward current and neuronal excitability are depressed with decreased O_2_ levels (10). In *Caenorhabditis elegans*, the hypoxia responses require cyclic guanosine monophosphate (cGMP)-gated cation channels (CNGs) including tax-2 and tax-4, and the atypical soluble guanylyl cyclases (sGCs) such as GCY-35 (9, 21, 76).

In *Drosophila*, oxygen sensing and behavioral responses to hypoxia were undescribed until a class of sGCs was identified, including Gyc88E, Gyc89Da and Gyc89Db (45) which contribute to the hypoxia response in *Drosophila* larvae (66). One of the CNG channel family members, CNGA, regulates an escape response to hypoxia in *Drosophila* larvae (66). However, similar responses have not been attributed to the other CNG channel family members, such as CNG-like (CNGL), CG3536 and CG17922. Flies with down-regulation of cGMP-specific phosphodiesterase (PDE) recover locomotor ability rapidly from anoxia, suggesting the involvement of cGMP in the modulation of anoxia recovery speed in adult flies (73). In *Drosophila* larvae, reduced levels of cGMP in O_2_-sensitive neurons result in longer times to respond to hypoxia, indicating that cGMP also regulates the escape response to hypoxia in *Drosophila* larvae (66). cGMP has several downstream targets, such as CNG channels and cGMP-dependent protein kinase G (PKG). Flies with lower PKG activity show an increased time to the onset of anoxic coma and are more behaviorally resistant to anoxia and hypoxia (11, 62). However, whether CNG channels regulate the anoxia or hypoxia responses of adult flies is unknown. Under anoxia, neural activity and most physiological functions are shutdown (53), whereas under hypoxia, the CNS remains functional, indicating that it is feasible to investigate the role of CNG channels in maintaining neural function and regulating locomotor activity under hypoxia. To date, little attention has been focused on one of the CNG channel family members, the CNGL channel. What we know about the CNGL channel can be summarized as follows:

1. *CNGL* is detected in fly optic lobe, central brain and thoracic ganglia, shows similarity to the mammalian CNG channel α and β subunits, and is predicted to form heteromeric channels with similar sensitivity to cAMP and cGMP (44, 66);
2. the CNGL channel is somehow involved in visual orientation memory (34), however, it does not contribute to the hypoxia escape response in *Drosophila* larvae (66);
3. A *CNGL* homolog in Hawaiian crickets is a candidate gene underlying interspecific variation in centrally-generated song patterns, which contribute to the rapid evolution of reproductive barriers and speciation (74).

Considering that cGMP and PKG are involved in responses to hypoxia and that CNG channels contribute to the escape response to hypoxia in *Drosophila* larvae yet CNGL channels do not, we investigated the role of the CNGL channel in regulating the hypoxia response of adult flies. We hypothesized that mutation of *CNGL* would alter behavioral and electrophysiological responses to hypoxia and tested the hypothesis using a locomotor assay and comparing *CNGL* mutant flies and w1118 control flies under normoxia, under hypoxia, and during recovery. In addition, we examined the effects of hypoxia on locomotion in pan-neural or pan-glial *CNGL* knockdown flies. We measured extracellular K^+^ concentration in the brain before and during hypoxia to investigate possible changes specific to K^+^ rather than overall ion homeostasis. In addition, we used fly lines with overexpression of Gyc88E or mutation of *Pde1c* to examine the interaction between CNGL and cGMP in response to hypoxia.

## 2 Materials and Methods

### 2.1 Flies

Fly strains and their sources: w1118 (L. Seroude laboratory, Queen’s University, Canada); CNGL^MB01092^ (Bloomington Stock Center, BSC #22988); UAS-CNGL-RNAi (BSC #28684); UAS-Gyc88E (D. Morton laboratory, Oregon Health & Science University, USA); Pde1c^KG05572^ (BSC #13901); elav-Gal4 (BSC #8765); and repo-Gal4 (BSC #7415).

Male progeny (CNGL^MB01092^/y) were obtained by crossing female CNGL^MB01092^ flies with male w1118 flies. This was to reduce as much as possible any differences in the genetic background of CNGL^MB01092^ and w1118 flies. Male progeny (CNGL^MB01092^/y; Pde1c^KG05572^/+;) were obtained by crossing female CNGL^MB01092^ flies with male Pde1c^KG05572^ flies. Further synchronization of the genetic background was difficult due to the lack of an eye-color marker in the CNGL^MB01092^, however, in the Discussion we argue against attributing our results to differences in genetic background.

Up- or down-regulation of a gene was carried out using the Gal4/UAS binary expression system (Brand and Perrimon, 1993; Clemens et al., 2000; Duffy, 2002; Hammond et al., 2000). Male progeny (; elav-Gal4/+; UAS-CNGL-RNAi/+ and; repo-Gal4/+; UAS-CNGL-RNAi/+) were obtained by crossing female UAS-CNGL-RNAi flies with elav-Gal4 or repo-Gal4 male flies, respectively. CNGL^MB01092^ flies also contained a CNGL enhancer-trap line (CNGL-Gal4), therefore, male progeny (CNGL^MB01092^/y; UAS-Gyc88E/+;) were obtained by crossing female CNGL^MB01092^ flies with UAS-Gyc88E male flies.

Flies were raised on standard medium (0.01% molasses, 8.2% cornmeal, 3.4% killed yeast, 0.94% agar, 0.18% benzoic acid, 0.66% propionic acid) at room temperature 21–23°C, 60–70% humidity. A 12h/12 h light/dark cycle was provided by three light bulbs (Philips 13 W compact fluorescent energy saver) with lights on at 7 am and off at 7 pm. Male flies were collected within two days after eclosion and raised for at least three additional days before the experiments. 100% N_2_ gas was used to knock down flies during collection but flies were not exposed to 100% N_2_ for 3 days prior to testing. All experiments took place between 10 am and 4 pm on flies younger than 9 days old.

### 2.2 RNA extraction and RT-PCR

Total RNA from around 20 whole flies of both sexes at 4-7 days old was extracted using a PureLink RNA Mini Kit (Cat# 12183020, ThermoFisher Scientific, USA). Reverse transcription (RT) was performed using a GoScriptTM Reverse Transcription System (A5001, Promega, USA). Polymerase chain reaction (PCR) conditions were: 95 ℃ for 30 min for hot start, followed by denaturing at 95 ℃ for 30 seconds, annealing at 60 ℃ for 1 minute, extension at 72 ℃ for 2 minutes for 40 cycles, and a final extension at 72 ℃ for 10 minutes. The primers were: CNGL-RF (5’-TCGTTCATCAGCGAGCATCC-3’, 5’-GGTGGCAACGTTCCTCTTGA-3’); CNGL-RD, RE, RJ and RI (5’-GGAGAGCTTCGCGTTTCCTG-3’, 5’-GAGGATGAGGATGTCGGTGC-3’). Tubulin 84B was used as loading control with primers 5’-CATGGTCGACAACGAGGCTA-3’ and 5’-GCACCTTGGCCAAATCACCT-3’. PCR products were separated in 1 - 2 % agarose gel, stained with 0.5 µg/ml ethidium bromide and visualized with UV light. Images of DNA bands were captured with AlphaImager Image Analysis System (Alpha Innotech).

### 2.3 Locomotor assay

The locomotor assay has been described (72). Illumination for video-capture was provided with a white light box (Logan Portaview slide/transparency viewer with 15 Watt color-corrected fluorescent bulbs) and the reflection from white cardboard screens surrounding the set up. Individual flies were gently aspirated into circular arenas (1.27 cm diameter; 0.3 cm depth) that prevented flying but allowed their walking to be video-recorded for later analysis. A custom-written fly tracking software using Open Computer Vision 2.0 (OpenCV2.0) was used for computing fly positions (the center of mass) and calculating path length per minute. Gas (air or the hypoxia mixture) continuously flowed through the apparatus at 2 L/min. After 5 minutes of adaptation in the arena and 10 minutes of locomotion recorded under normoxia, hypoxia was induced by switching from air to a mixture of 2% O_2_ and 98% N_2_ for 15 minutes. These parameters for the hypoxia were chosen based on preliminary results indicating a good balance between the severity of the effect and the ability to recover in a short enough time to prevent the flies being stressed in the locomotor chambers. The ratio of O_2_ and N_2_ chosen here was based on previous studies, which define such a composition as “hypoxia” (2, 15). After switching back to air, the flies recovered under normoxia for 40 min. Experiments were conducted in the light and at least three hours away from light-dark transit, in order to avoid morning and evening activity peaks (22). Flies do not walk continuously in one direction in the arena but pause for variable periods and/or change direction (71). To estimate the frequency of pausing during the first 10 min in normoxia, we counted the number of 1 second intervals during which the path length was zero (e.g. a fly motionless for 10 min would have 600 seconds of zero path length, represented as 600 stops/10 min). To estimate maximum walking speed, we chose the farthest distance travelled in 1 second during the first 10 min, assuming that during normoxia each fly would have at least one 1 second episode of continuous walking. Every minute the total path length, including stationary periods, was averaged and plotted under three different conditions including normoxia (10 min), hypoxia (15 min), and recovery after hypoxia (40 min). In addition, the path length per minute under normoxia was averaged over 10 minutes (1-10 min) (L_normoxia_), under hypoxia during the last 10 minutes (16-25 min) (L_hypoxia)_ and under normoxia during the first 10 minutes after 5 minutes adaptation (31-40 min) and the last 10 minutes of recovery (56-65 min) (L_recovery_). L_normoxia_ was compared within and between fly genotypes. To determine the effect of O_2_ level changes on fly locomotion, L_normoxia_, L_hypoxia_ and L_recovery_ were compared within the same fly line. Percent recovery was calculated only for the fly lines with significant effects of O_2_ level changes on fly locomotion (*P* < 0.05 when performing Kruskal-Wallis test). For each fly, the percent recovery after hypoxia was compared within and between fly genotypes using the values for L_normoxia_ and L_recovery_ (i.e. the final 10 minutes of recovery).

To investigate the effect of hypoxia on flies with *CNGL* down-regulation or knockdown, the path length during the last 5 minutes of normoxia, 15 minutes hypoxia and the first 5 minutes recovery after hypoxia (6-30 min in total) was plotted again in the same figure. During hypoxia, path length per minute quickly reached an acute increase in path length per minute and then sharply dropped, followed by a gradual increase. The peak value of this acute increase (11 or 12 min) was compared with the last minute prior to hypoxia (10 min).

### 2.4 Recording of CNS extracellular potassium ion concentrations

CNS extracellular potassium ion concentration ([K^+^]_o_) was measured with a K^+^-sensitive microelectrode. K^+^-sensitive microelectrodes were prepared as previously described (3). Non-filamented glass capillaries (1B100-4, World Precision Instruments, USA) were immersed into 99.9 % methanol for five minutes and dried out at room temperature. Low resistance electrodes (5-10 megaohms) were fashioned from cleaned capillaries using the P-87 Faming/Brown micropipette puller (Sutter Instrument Co.). The electrodes were silanized with dichlorodimethylsilane (440272-100ml, Sigma-Aldrich) on a hot plate (100 °C) for 1 hour. Silanization was repeated once. After cooling, the electrode tip was filled with potassium ionophore I - cocktail B (99373, Sigma-Aldrich) for a distance of 200-1000 µm. Electrodes were then backfilled with 1 M KCl. The electrode tips were immersed in distilled water until use. K^+^-sensitive microelectrodes were freshly prepared and discarded after 12 hrs. Reference electrodes (5-10 megaohms) were pulled using filamented capillaries (1B100F-4, World Precision Instruments, USA) and back-filled with 1 M KCl. Electrical signals were acquired using AxoScope 10 software (Molecular Devices) with a pH/ION amplifier (Model 2000, A-M Systems) and a digitizer (Digidata 1550A, Molecular Devices). The K^+^-sensitive electrode was calibrated with two KCl solutions (10 and 100 mM) at room temperature. Electrodes were kept for experiments if there was a voltage change of 50 - 58 mV corresponding to a 10-fold change in K^+^ concentrations. The voltage output of the K^+^-sensitive electrode was set to zero in 10 mM KCl.

A fly was secured in a trimmed pipette tip (200 µl) with its head exposed and thorax and abdomen inside. A drop of wax was placed underneath the head to limit movement. A small area of cuticle between the compound eyes was removed. Both K^+^-sensitive and reference electrodes were inserted into the middle of the brain. A grounding wire (Ag/AgCl wire) was inserted into the thorax. Flies recovered from experimental handling for 5 minutes prior to recording. Hypoxia was delivered using a mixture of 2 % O_2_ and 98 % N_2_ at 5 L/min for 1 minute, followed by recovery in air flow for 5 minutes. The hypoxia duration was shorter than for the locomotor assays because the fly preparations were harsher and the recordings more difficult to maintain. Moreover, in these experiments we were interested in the recovery of [K^+^]_o_ whereas recovery of locomotion involves restoration of multiple additional processes and takes longer. Recordings were made in the daytime under regular light illumination.

K^+^-sensitive electrodes could be impaired during the insertion into the tissue. Therefore, a second calibration was performed after the recording by placing the ion-selective electrode into 10 mM KCl. A difference of < 5 mV between calibrations was acceptable, otherwise the recording was discarded. Conversion from voltage to extracellular K^+^ concentration was performed using the Nernst equation: [K^+^]_o_ = 10 ×10^voltage/58.2^.

### 2.5 Analysis of variables in response to hypoxia

The variables measured during hypoxia included: 1) extracellular K^+^ concentration change obtained by the K^+^-sensitive electrode (Δ [K^+^]_o_); 2) time to reach peak of [K^+^]_o_ (t_peak_); and 3) time to recovery (t_recovery_) (see Figure 8A).

The baselines in [K^+^]_o_ trace were obtained one minute before hypoxia treatment. The parameter Δ [K^+^]_o_ was defined as the concentration difference between the peak and the baseline. The parameter t_peak_ was defined as the time required to reach the peak from the onset of the hypoxia treatment. During recovery, [K^+^]_o_ dropped quickly and stabilized within 3.5 min, therefore, the last 1 min data during the 5 min recovery period was chosen to calculate the final baseline after hypoxia. The variable t_recovery_ was defined as the first time to reach the final baseline after hypoxia was turned off.

### 2.6 Statistics

Three replicates of each experiment were conducted. The sample sizes for each test are stated in the Results section. It should be noted that the sample sizes for the experiments varied depending on the fly availability, due to the inconsistent yields of progeny obtained after fly crosses. Statistical analysis was performed using Prism version 5.0 (GraphPad Software, San Diego, CA). A D’Agostino & Pearson omnibus normality test was conducted to examine the data distribution. Because some of the data had non-Gaussian distributions, nonparametric Mann-Whitney or Kruskal-Wallis tests with post-hoc comparisons were performed to examine the difference of medians between groups. Kruskal-Wallis test (without post-hoc comparisons) was performed to determine the effect of O_2_ level changes within the same fly line. *P* < 0.05 was considered as indicating statistical significance.

## 3 Results

### 3.1 *CNGL* transcriptional down-regulation in CNGL^MB01092^ flies

RT-PCR was used to examine transcriptional changes in *CNGL* mutants. According to the National Center for Biotechnology Information (NCBI) nucleotide database, there are five mRNA variants including CNGL-RD, RE, RJ, RI and RF for *CNGL* (Figure 1A). Among them, CNGL^MB01092^ showed a reduction of CNGL-RD, RE, RJ and RI transcripts compared with w1118 (Figure 1B). The other variant, CNGL-RF, was likely unchanged. Thus, CNGL^MB01092^ flies had reduced levels of *CNGL* transcripts.

**Figure 1.**
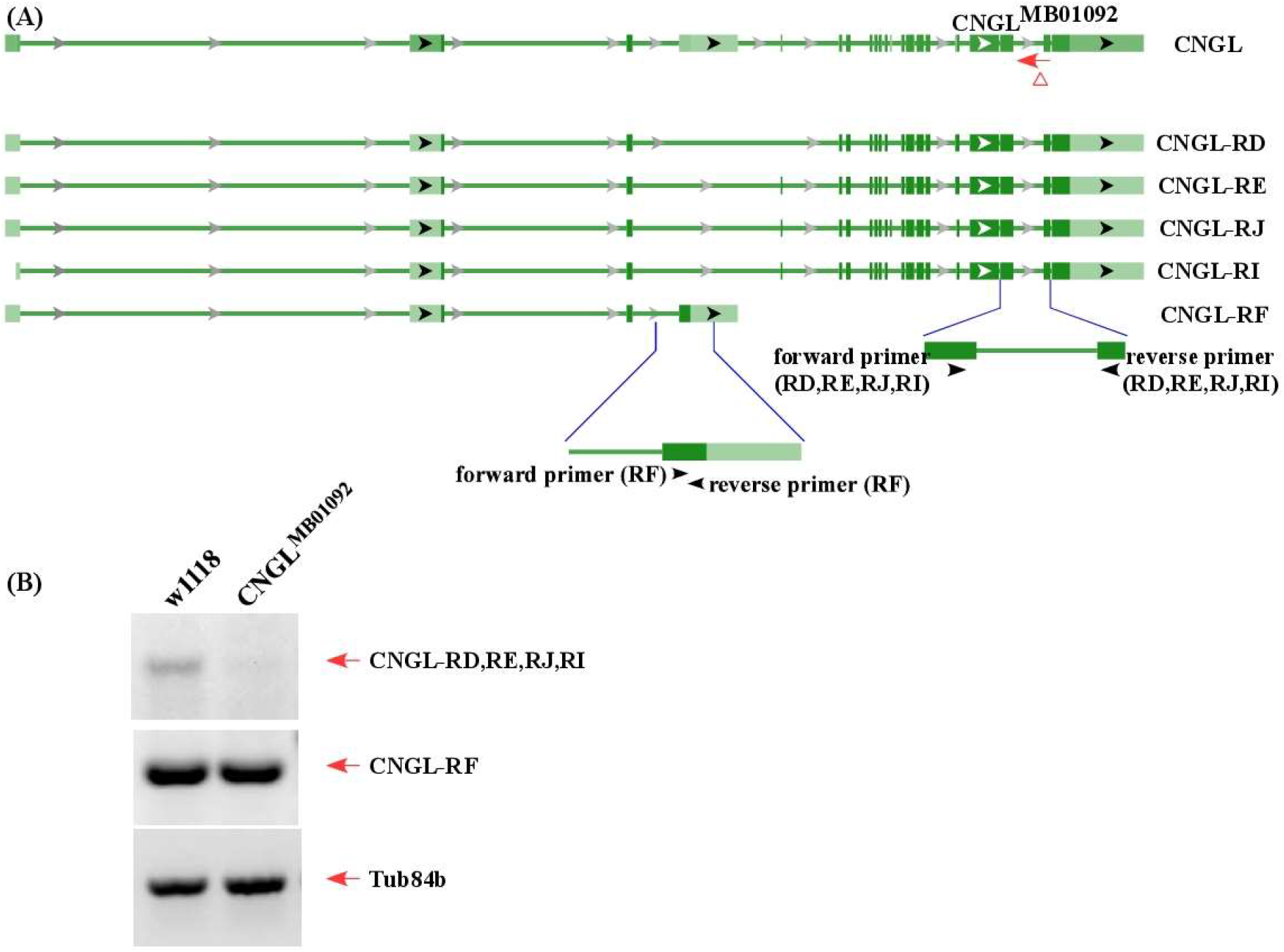
Low level of *CNGL* transcripts in CNGL^MB01092^. (A) Five predicted transcripts of *CNGL* (CNGL-RD, RE, RJ, RI and RF), two sets of primers and CNGL^MB01092^ mutant line. (B) RT-PCR was performed on total RNA extracted from whole flies of either w1118 or CNGL^MB01092^. For a loading control, the cDNA was also amplified with Tub84b primers. Lower level of *CNGL* transcripts was observed in the mutant.

### 3.2 Reduced locomotion under normoxia and reduced recovery from hypoxia in CNGL^MB01092^/y flies

Locomotor activity was analyzed under normoxia, under hypoxia, and during recovery. During the 10 min period of normoxia, w1118 flies walked farther in each minute than CNGL^MB01092^/y flies (Figure 2A-C). Additionally, w1118 flies had significantly fewer stops of at least 1 second (n=14, median 83 stops/10 min, interquartile range (IQR) 25.3-108.8 stops/10 min) than CNGL^MB01092^/y flies (n=12, median 259.5 stops/10 min, IQR 170.3-368.5 stops/10 min) (*P* < 0.001, Mann-Whitney test). w1118 flies also walked faster (maximum speed: n=14, median 1.3 cm/s, IQR 1.2-1.5 cm/s) than CNGL^MB01092^/y flies (maximum speed: n=12, median 0.8 cm/s, IQR 0.6-1.1 cm/s) (*P* < 0.001, Mann-Whitney test). Under hypoxia, the path length per minute of w1118 flies dropped, however, in CNGL^MB01092^/y flies it showed an acute increase within 1 minute before dropping, which was followed by a gradual increase (Figure 2A-C, see also 7A). During the recovery period, *CNGL* mutants showed almost no activity, while w1118 flies had a larger path length per minute (Figure 2 A-C).

**Figure 2.**
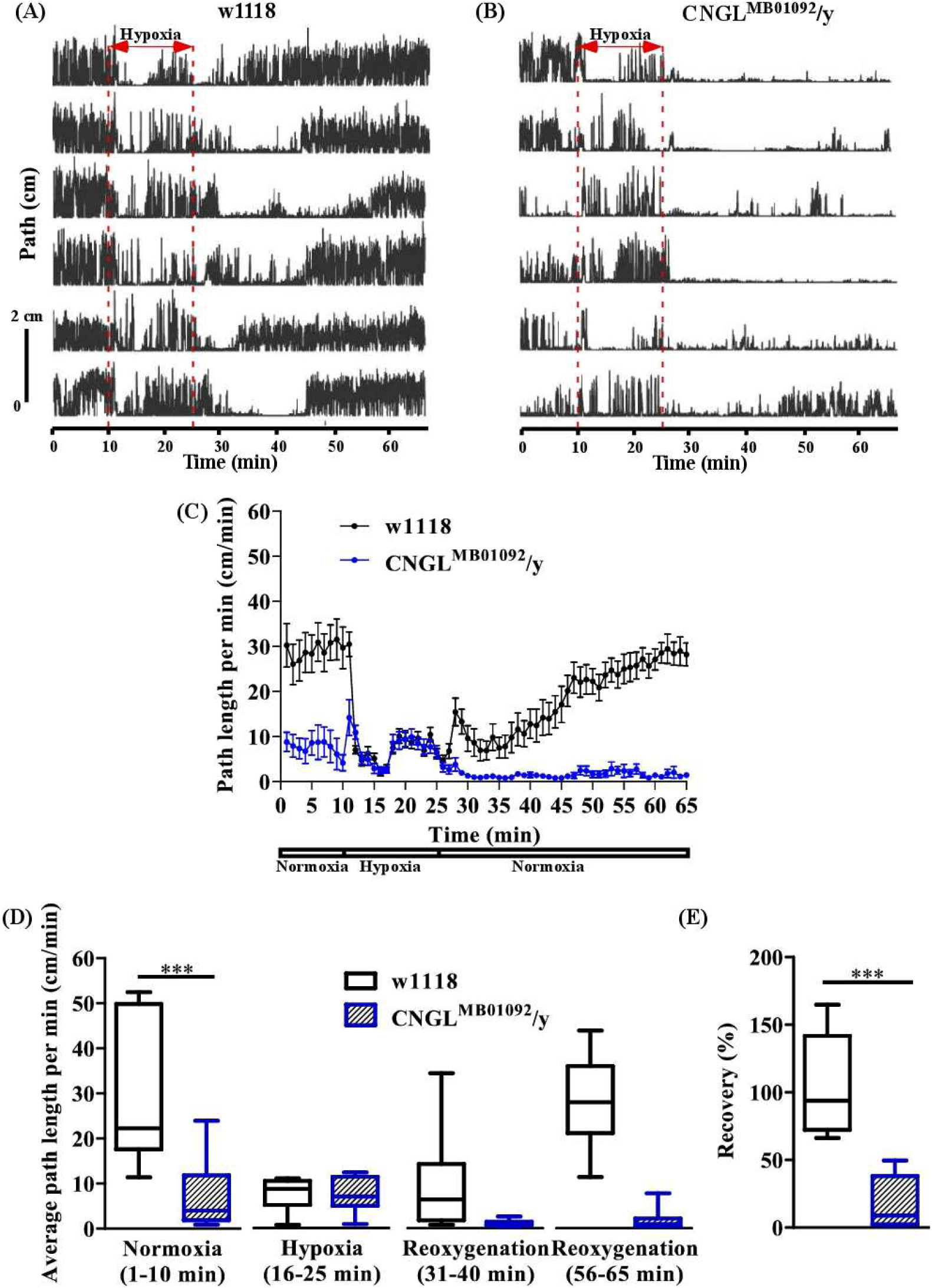
Reduced locomotion under normoxia and reduced recovery in response to hypoxia in CNGL^MB01092^/y flies. (A-B) Locomotor analysis for w1118 flies and CNGL^MB01092^/y flies with hypoxia. Each plot represents locomotor activity of a single fly. The positions of individual fly were measured every 0.2 s. A 15 min hypoxia was applied after 10 min locomotion under normoxia. Flies were then returned to normoxia and allowed to recover for 40 min. (C) Path length per minute of control flies (w1118) (black circles and black line, n=14) and mutant fly line CNGL^MB01092^/y (blue circles and blue line, n=12) under normoxia, during hypoxia and after hypoxia-treatment. (D) The average path length per minute under normoxia (1-10 min), during hypoxia (16-25 min), and during reoxygenation (31-40 min and 56-65 min) of w1118 and CNGL^MB01092^/y flies. (E) The percent recovery of w1118 and CNGL^MB01092^/y flies in response to hypoxia. Asterisks (***) indicate *P* < 0.001 by Mann-Whitney test.

Under normoxia, w1118 flies walked farther (n=14, median 22.3 cm/min, IQR 17.6-49.8 cm/min) than CNGL^MB01092^/y flies (n=12, median 4.0 cm/min, IQR 1.9-11.8 cm/min) (*P* < 0.001, Mann-Whitney test) (Figure 2D, left panel). Therefore, the *CNGL* mutation obtained from CNGL^MB01092^/y flies was associated with reduced locomotor activity under normoxia. Alterations of O_2_ level had a significant influence on fly locomotion in w1118 flies (*P* < 0.001, Kruskal-Wallis test without post-hoc test) (Figure 2D). Similarly, locomotion in CNGL^MB01092^/y flies also varied with the O_2_ level changes (*P* < 0.001, Kruskal-Wallis test) (Figure 2D). CNGL^MB01092^/y flies showed a reduced recovery (n=12, median 8.9%, IQR 2.3-38.1%) compared with w1118 control flies (n=14, median 93.9%, IQR 72.3-141.7%) (*P* < 0.001, Mann-Whitney test) (Figure 2E). Thus, the mutation in CNGL^MB01092^/y flies led to reduced recovery from hypoxia.

### 3.3 Reduced locomotion under normoxia and reduced recovery in response to hypoxia in flies with *CNGL* knockdown in neurons

To examine the relationship between reduced locomotion under normoxia and fly genotypes, we first targeted RNAi knockdown of *CNGL* to the central nervous system with the pan-neuronal driver elav-Gal4 (3, 14, 40). No apparent morphological abnormality or developmental delay was observed in the *CNGL* pan-neuronal knockdown flies compared with controls.

Under normoxia, the locomotor performance of;elav-Gal4/+; UAS-CNGL-RNAi/+ males was lower than controls (Figure 3A-D). The average path length per minute of the mutant flies (n=11, median 2.6 cm/min, IQR 1.2-13.8 cm/min) was severely reduced compared with controls (;elav-Gal4/+;, n=8, median 35.5 cm/min, IQR 32.2-39.2 cm/min, and;;UAS-CNGL-RNAi/+, n=18, median 38.6 cm/min, IQR 34.5-41.1 cm/min) (*P* < 0.01 or 0.001, respectively, Kruskal-Wallis test with Dunn’s multiple comparison) (Figure 3E, left panel). Similar to the CNGL^MB01092^/y flies, flies with *CNGL* knockdown in neurons also showed an acute increase in path length per minute at the beginning of hypoxia (Figure 3C-D and 7B). The O_2_ level changes had a significant effect on fly locomotor activities in all the three fly genotypes (*P* < 0.001 for all three comparisons within same genotype, Kruskal-Wallis test without post-hoc test) (Figure 3E). During the recovery after hypoxia,;elav-Gal4/+; UAS-CNGL-RNAi/+ males showed reduced locomotor performance compared with controls (Figure 3A-D). The percent recovery of mutant flies (n=11, median 6.1%, IQR 4.4-9.6%) was significantly lower than controls (;elav-Gal4/+;, n=8, median 57.5%, IQR 31.3-69.8%, and;;UAS-CNGL-RNAi/+, n=18, median 62.1%, IQR 59.7-65.9%) (*P* < 0.01 or 0.001, respectively, Kruskal-Wallis test with Dunn’s multiple comparison) (Figure 3F).

**Figure 3.**
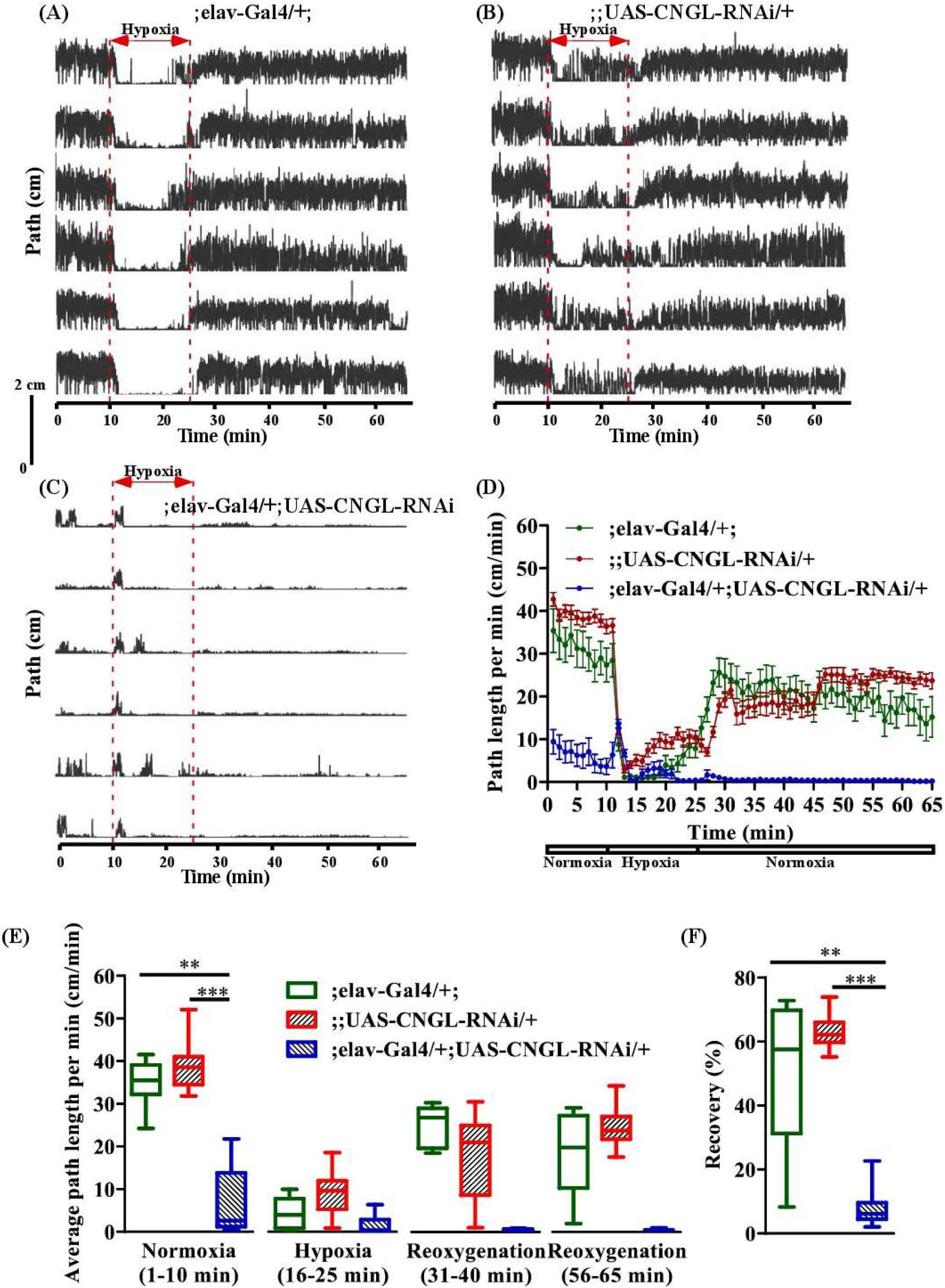
Reduced locomotion under normoxia and reduced recovery in response to hypoxia in flies with *CNGL* knockdown in neurons. (A-C) Locomotor analysis for flies with *CNGL* knockdown pan-neuronally and control flies with hypoxia. (D) Path length per minute of control flies (;elav-Gal4/+;, green circles and green line, n=8, and;;UAS-CNGL-RNAi/+, red circles and red line, n=18) and *CNGL* pan-neuronal knockdown flies (; elav-Gal4/+; UAS-CNGL-RNAi/+, blue circles and blue line, n=11) during normoxia, hypoxia and recovery process. (E) The average path length per minute under normoxia (1-10 min), during hypoxia (16-25 min), and during reoxygenation (31-40 min and 56-65 min) of control flies and flies with *CNGL* knockdown in neurons. (F) The percent recovery of control and *CNGL* pan-neuronal knockdown flies. Asterisks (** or ***) indicate *P* < 0.01 or 0.001, respectively, by Kruskal-Wallis test with Dunn’s multiple comparison.

### 3.4 Reduced locomotion under normoxia in flies with *CNGL* knockdown in glia

The RNAi knockdown of *CNGL* using the pan-glial driver repo-Gal4 (58) was targeted as well. Similarly, no apparent morphological abnormality or developmental delay was observed.

The RNAi knockdown of *CNGL* with the pan-glial driver led to greatly reduced locomotor performance compared with controls under normoxia (Figure 4A-D).;;repo-Gal4/UAS-CNGL-RNAi males displayed significantly lower average path length per minute (n=8, median 0.8 cm/min, IQR 0.4-1.3 cm/min) compared with controls (;;repo-Gal4/+, n=10, median 48.1 cm/min, IQR 43.1-55.7 cm/min, and;;UAS-CNGL-RNAi/+, n=12, median 38.8 cm/min, IQR 37.7-44.4 cm/min) (*P* < 0.001 or 0.05, respectively, Kruskal-Wallis test with Dunn’s multiple comparison) (Figure 4E, left panel). The changes in O_2_ level affected locomotion only in the two control flies (;;repo-Gal4/+ and;;UAS-CNGL-RNAi/+, both *P* < 0.001, Kruskal-Wallis test without post-hoc test), however, the minimal locomotor activity in;;repo-Gal4/UAS-CNGL-RNAi flies did not vary when O_2_ level was altered (*P* > 0.05, Kruskal-Wallis test without post-hoc test) (Figure 4E). An acute increase in path length per minute representing uncoordinated movement could be observed in;;repo-Gal4/UAS-CNGL-RNAi flies at the beginning of hypoxia treatment (Figure 4C-D and 7C).

**Figure 4.**
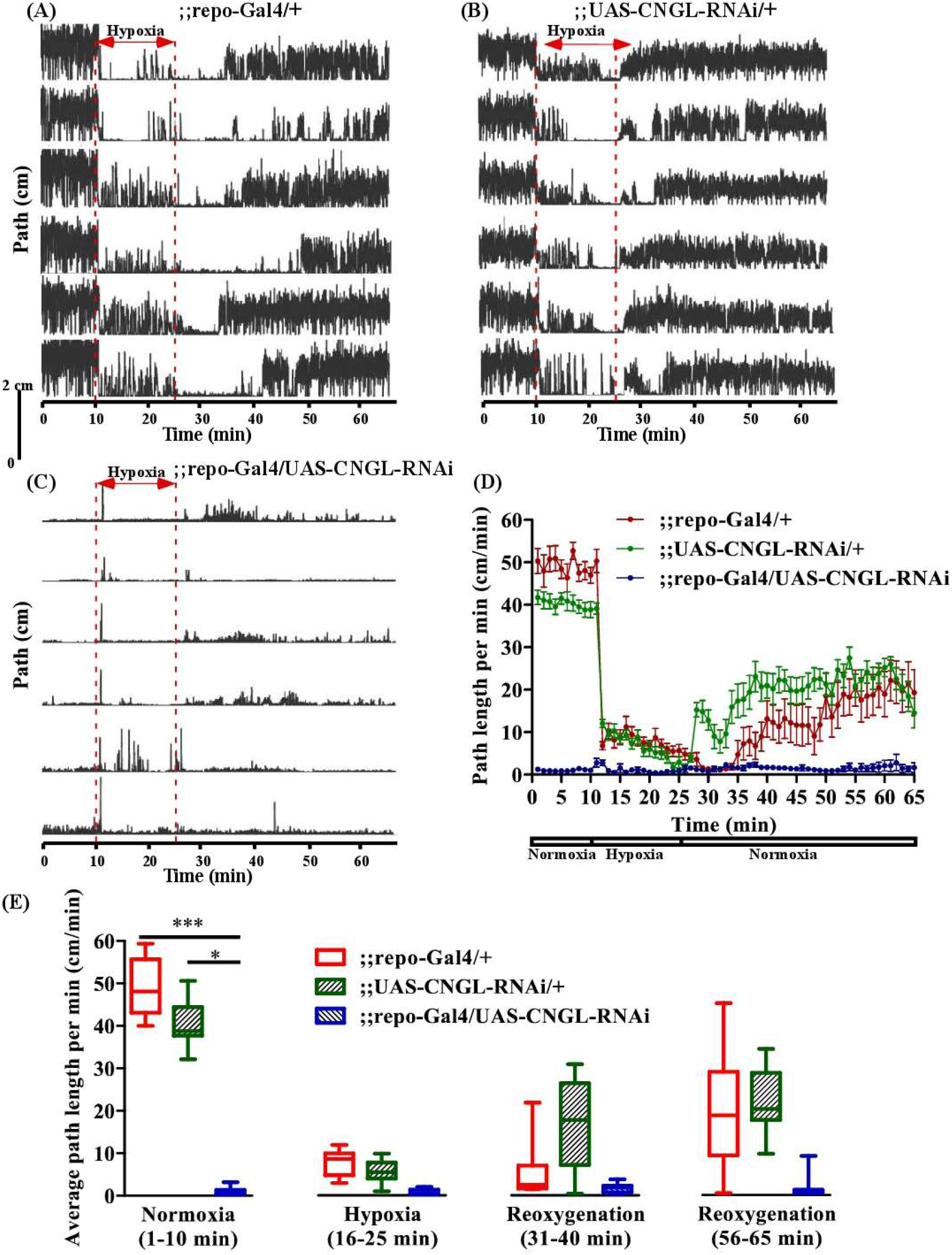
Reduced locomotion under normoxia in flies with *CNGL* knockdown in glia. (A-C) Locomotor analysis for flies with pan-glial *CNGL* down-regulation and control flies with hypoxia. (D) Path length per minute of control flies (;;repo-Gal4/+;, red circles and red line, n=10, and;;UAS-CNGL-RNAi/+, green circles and green line, n=12) and *CNGL* pan-glial knockdown flies (;;repo-Gal4/UAS-CNGL-RNAi, blue circles and blue line, n=8) during normoxia, hypoxia and recovery process. (E) The average path length per minute under normoxia (1-10 min), during hypoxia (16-25 min), and during reoxygenation (31-40 min and 56-65 min) of control flies and flies with *CNGL* knockdown in glia. Asterisks (* or ***) indicate *P* < 0.05 or 0.001, respectively, by Kruskal-Wallis test with Dunn’s multiple comparison.

### 3.5 Overexpression of Gyc88E eliminated the reduced recovery in response to hypoxia in *CNGL* mutant

cGMP regulates the behavioral responses to hypoxia in *Drosophila* larvae (66), and modulates the onset of anoxic coma (11) and the speed of anoxic recovery in adult flies (73). However, it is unclear whether cGMP modulates locomotor activity under normoxia and whether it regulates the hypoxia response in adult flies. Moreover, the *CNGL* mutants were not lacking all transcripts and it would be possible for increased activation of a reduced number of channels to mitigate the locomotor impairment. Therefore, an atypical soluble guanylyl cyclase Gyc88E, which catalyzes cGMP production, was expressed in *CNGL* mutant flies to examine whether cGMP upregulation could compensate for the effects of *CNGL* downregulation.

Under normoxia, the flies CNGL^MB01092^/y; UAS-Gyc88E/+; had better locomotor performance compared with CNGL^MB01092^/y flies (Figure 5A-D). Although the average path length per minute of flies CNGL^MB01092^/y; UAS-Gyc88E/+; (n=16, median 43.2 cm/min, IQR 39.5-49.5 cm/min) was comparable to flies;UAS-Gyc88E/+; (n=25, median 44.5 cm/min, IQR 36.9-48.9 cm/min) (*P* > 0.05, Kruskal-Wallis test with Dunn’s multiple comparison), the average speed of flies CNGL^MB01092^/y; UAS-Gyc88E/+; was significantly higher compared with CNGL^MB01092^/y flies (n=14, median 6.2 cm/min, IQR 3.2-10.2 cm/min) (*P* < 0.001, Kruskal-Wallis test with Dunn’s multiple comparison) (Figure 5E, left panel). The alterations in O_2_ level affected locomotion in all three fly lines (*P* < 0.001 for all three comparisons within same genotype, Kruskal-Wallis test without post-hoc test) (Figure 5E). The CNGL^MB01092^/y flies also showed an acute increase in path length per minute within 2 min at the onset of hypoxia (Figure5A and D, and 7D).

**Figure 5.**
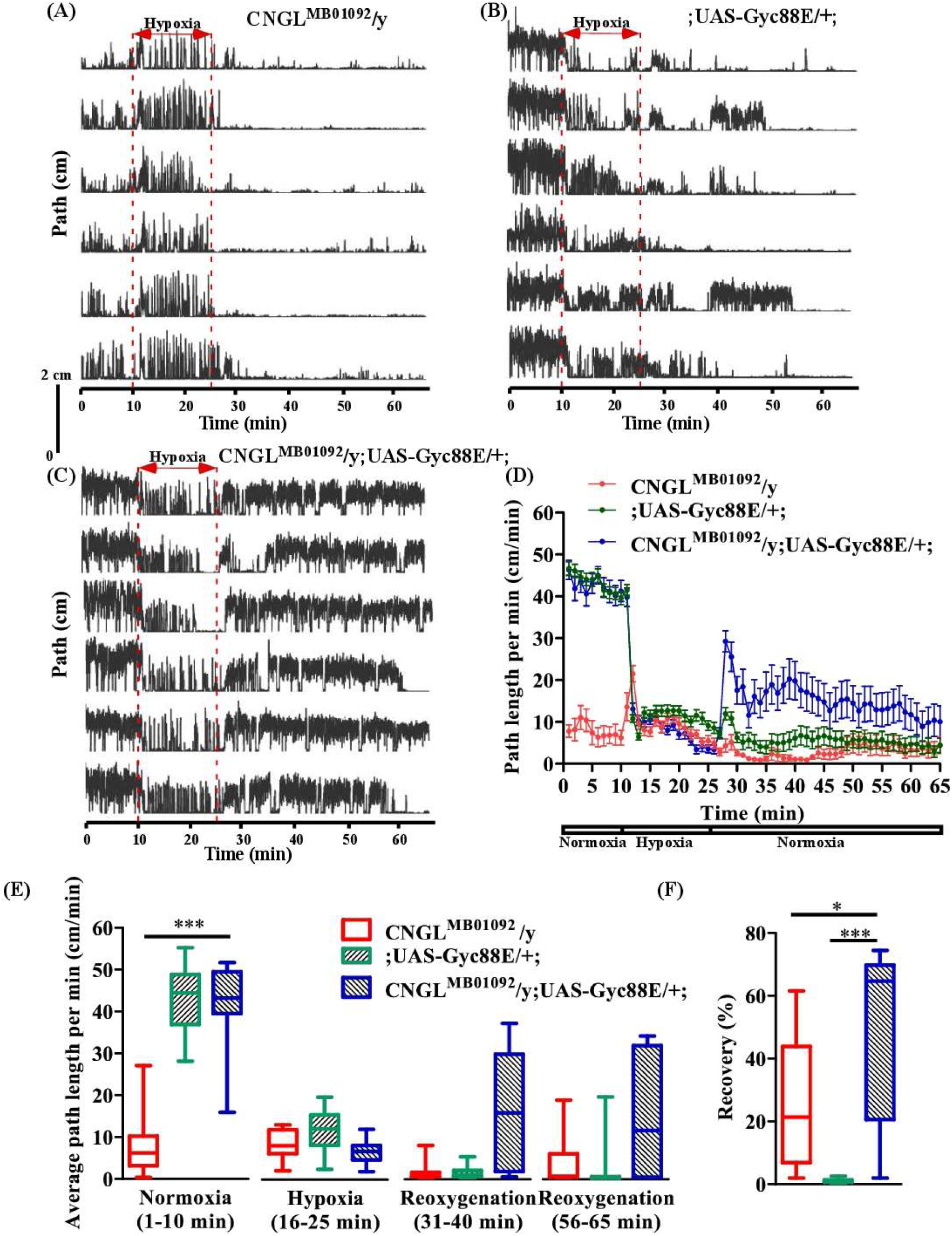
Overexpression of Gyc88E eliminated both the reduced locomotion under normoxia and the reduced recovery in response to hypoxia in CNGL mutant. (A-C) Locomotor analysis for flies with Gyc88E overexpression and control flies with hypoxia. (D) Path length per minute of control flies (CNGL^MB01092^/y, red circles and red line, n=12, and;UAS-Gyc88E/+;, green circles and green line, n=29) and *CNGL* mutant flies with Gyc88E overexpression (CNGL^MB01092^/y; UAS-Gyc88E/+;, blue circles and blue line, n=16) during normoxia, hypoxia and recovery. (B) The average path length per minute under normoxia (1-10 min), during hypoxia (16-25 min), and during reoxygenation (31-40 min and 56-65 min) of control flies and *CNGL* mutant flies with Gyc88E overexpression. (C) The percent recovery of control flies and *CNGL* mutant flies with Gyc88E overexpression. Asterisks (* or ***) indicate *P* < 0.05 or 0.001, respectively, by Kruskal-Wallis test with Dunn’s multiple comparison.

During recovery, CNGL^MB01092^/y; UAS-Gyc88E/+; flies walked farther compared with the two control fly lines (Figure 5A-D). Overexpression of Gyc88E in *CNGL* mutant resulted in larger percent recovery (n=16, median 64.7%, IQR 20.6-69.8%) while the two control fly lines, CNGL^MB01092^/y and;UAS-Gyc88E/+;, had percentage recoveries of 21.3% (n=14, IQR 6.9-43.8%), and 1.0% (n=25, IQR 0.7-1.4%), respectively (*P* < 0.05 or 0.001, respectively, Kruskal-Wallis test with Dunn’s multiple comparison) (Figure 5F).

Therefore, the overexpression of Gyc88E in *CNGL* mutant flies suppressed the reduced recovery in response to hypoxia.

### 3.6 *Pde1c* mutation eliminated the reduced recovery in response to hypoxia in*CNGL* mutant

Overexpression of Gyc88E, which leads to the upregulation of cGMP suppresses the *w*-RNAi induced delay of locomotor recovery (73) and eliminates the reduced recovery in response to hypoxia in *CNGL* mutant. Genes for cGMP-specific PDEs, which regulate intracellular levels of cGMP by hydrolyzing cGMP, have been identified in *Drosophila* (12, 47). Therefore, *PDE* mutation is another approach to investigate the impact of cGMP upregulation (73). One of the *PDE* mutants, *Pde1c*^KG05572^, could compensate for the *w*-RNAi-induced delay of locomotor recovery from anoxia. Here we examined whether the *Pde1c* mutation could also eliminate the reduced locomotion under normoxia and the reduced recovery in response to hypoxia in *CNGL* mutant flies.

Among the three fly genotypes under normoxia, flies CNGL^MB01092^/y; *Pde1c*^KG05572^/+; displayed better locomotor performance compared with *CNGL* mutant flies (Figure 6A-D). The average path length per minute in CNGL^MB01092^/y; *Pde1c*^KG05572^*/+*; was 31.2 cm/min (n=15, IQR 29.2-34.0 cm/min). Even though it showed smaller path length per minute than flies; *Pde1c*^KG05572^/+; (n=12, median 48.2 cm/min, IQR 42.8-51.7 cm/min), it had larger value than *CNGL* mutant flies CNGL^MB01092^/y flies (n=8, median 7.3 cm/min, IQR 2.4-18.5 cm/min) (*P* < 0.001 for both comparisons, Kruskal-Wallis test with Dunn’s multiple comparison) (Figure 6E, left panel). These data indicate that *Pde1c* mutation suppressed the reduced locomotor activity under normoxia in *CNGL* mutant. In addition, O_2_ level changes affected fly locomotor activities in all three fly genotypes (*P* < 0.001 for all three comparisons within same genotypes, Kruskal-Wallis test without post-hoc test) (Figure 6E). Similarly, the path length per minute in CNGL^MB01092^/y flies reached to a peak value at the beginning of the hypoxia treatment and immediately dropped ~10 cm/min, which was then followed by a gradual increase (Figure 6A and D, and 7E).

**Figure 6.**
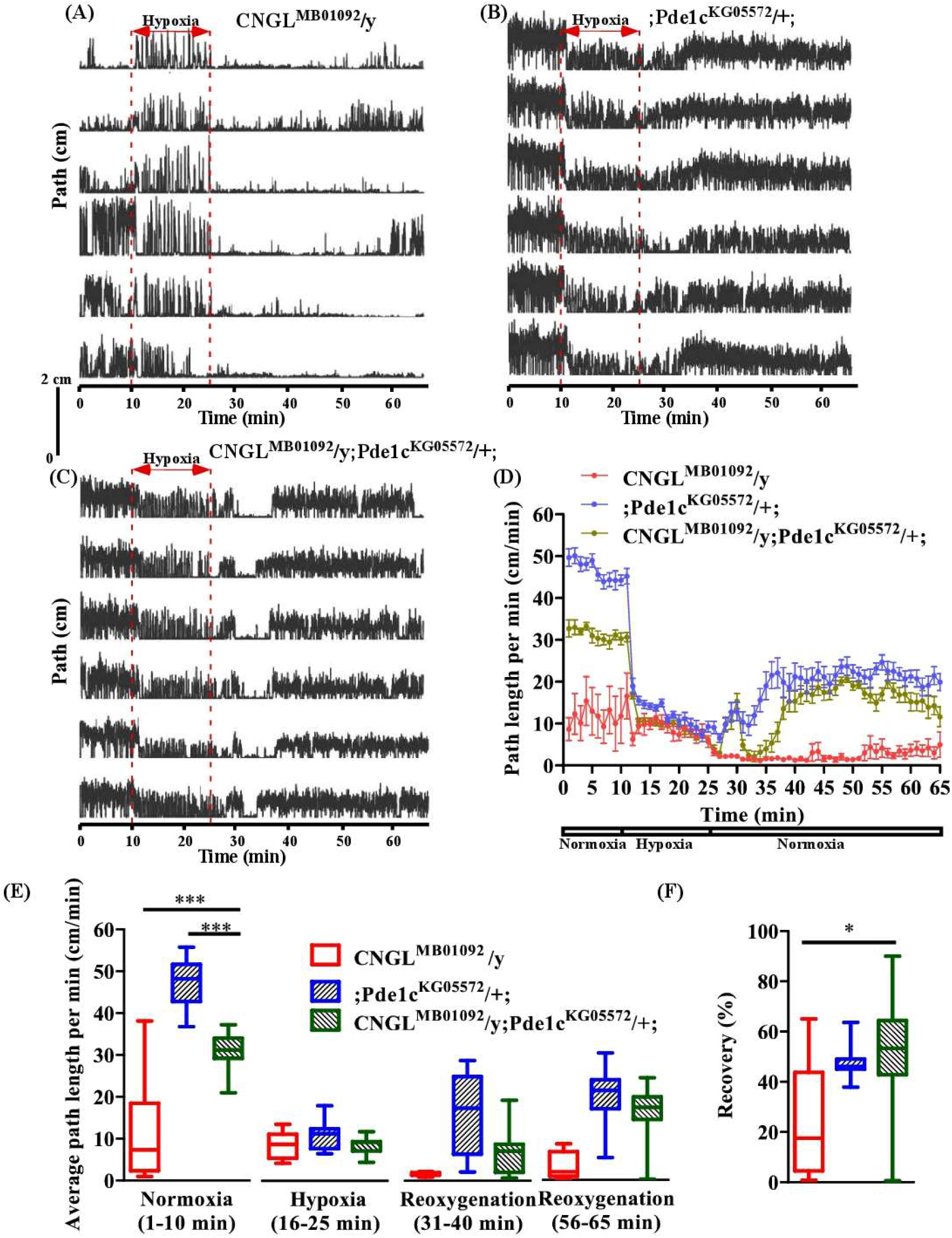
*Pde1c* mutation eliminated both the reduced locomotion under normoxia and the reduced recovery in response to hypoxia in *CNGL* mutant. (A-C) Locomotor analysis for flies with *Pde1c* mutation and control flies with hypoxia. (D) Path length per minute of control flies (CNGL^MB01092^/y, red circles and red line, n=8, and;Pde1c^KG05572^/+;, blue circles and blue line, n=12) and *CNGL* mutant flies with *Pde1c* mutation (CNGL^MB01092^/y; *Pde1c*^KG05572^/+;, green circles and green line, n=15) during normoxia, hypoxia and recovery. (B) The average path length per minute under normoxia (1-10 min), during hypoxia (16-25 min), and during reoxygenation (31-40 min and 56-65 min) of control flies and *CNGL* mutant flies with *Pde1c* mutation. (C) The percent recovery of control flies and *CNGL* mutant flies with *Pde1c* mutation. Asterisks (* or ***) indicate *P* < 0.05 or 0.001, respectively, by Kruskal-Wallis test with Dunn’s multiple comparison.

During recovery, flies CNGL^MB01092^/y; *Pde1c*^KG05572^/+; walked farther relative to CNGL^MB01092^/y flies (Figure 6A-D). Although the recovery percentage in CNGL^MB01092^/y; *Pde1c*^KG05572^*/+*; (n=15, median 53.3%, IQR 42.8-64.4%) was comparable with that of; *Pde1c*^KG05572^*/+*; flies (n=12, median 46.0%, IQR 45.1-49.0%), the value of CNGL^MB01092^/y; *Pde1c*^KG05572^*/+*; was significantly higher than that of CNGL^MB01092^/y flies (n=8, median 17.5%, IQR 4.6-43.8%) (*P* > 0.05 or < 0.05, respectively, Kruskal-Wallis test with Dunn’s multiple comparison) (Figure 6F). Therefore, the result suggests that the mutation in *Pde1c* eliminated the reduced recovery percentage in response to hypoxia in *CNGL* mutant flies.

### 3.7 Increased locomotion at the onset of hypoxia in flies with *CNGL* down-regulation or pan-neuronal knockdown

Flies with *CNGL* down-regulation or knockdown showed increased locomotion within 1-2 min at the onset of hypoxia and the path length per minute then dropped rapidly; afterwards, the value gradually increased (Figure 7A-E, left panel (summarized from Figure 2D-6D)). The peak value of this acute increase in path length per minute was then compared with that at 10 min, which was the last minute before hypoxia. In most cases, the peak value was significantly higher than that at 10 min (Figure 6A, right panel: *P* < 0.05; Figure 6B, right panel: *P* < 0.01; Figure 6D, right panel: *P* < 0.001; Mann-Whitney test for all comparisons). However, no significant differences were observed in flies with *CNGL* knockdown in glia (Figure 6C, right panel: *P* > 0.05, Mann-Whitney test). Although an acute increase was observable in Figure 7E, the value was not remarkably higher than that at 10 min, which might be due to large variations (*P* > 0.05, Mann-Whitney test).

**Figure 7.**
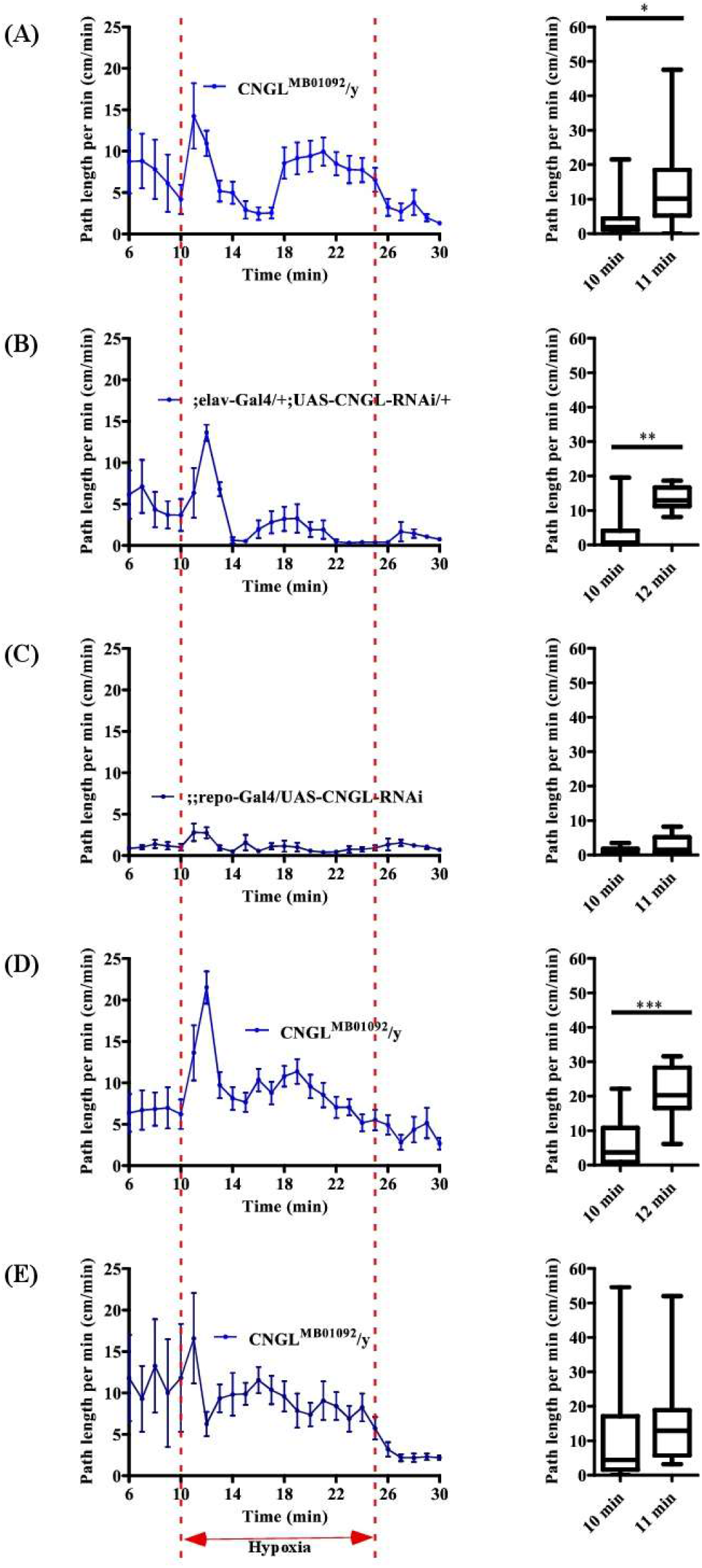
Increased locomotion at the onset of hypoxia in flies with *CNGL* down-regulation or pan-neuronal knockdown. Left panels of A-E are summarized from Figure 4.2D-4.6D, respectively. Right panels of A-E are peak value of the acute increase in path length per minute and at 10 min in flies with *CNGL* down-regulation or knockdown.

### 3.8 Reduced basal levels of CNS extracellular K^+^ in CNGL^MB01092^/y flies

In vertebrate photoreceptors, CNG channels determine the dynamics of Ca^2+^ homeostasis, which plays a key role in light adaptation (31). Additionally, in olfactory receptor neurons, CNG channels are permeable to Ca^2+^ ions, which contributes to feedforward and feedback mechanisms (30). CNGL channels are strongly expressed in the brain and thoracic ganglia (44) and have structural similarity with hyperpolarization-activated cyclic nucleotide-gated HCN channels known to be involved in central pattern generation (74). This background suggests that mutations of CNG channels, such as the *CNGL* mutation, could affect extracellular ion homeostasis in the brain.

The K^+^-selective and reference electrodes were inserted into the middle of the brain to measure [K^+^]_o_. After 5 minutes adaptation to the recording system, [K^+^]_o_ baseline was stable throughout the experiment. The *CNGL* mutation in CNGL^MB01092^/y flies (n=19, median 6.2 mM, IQR 5.4-6.8 mM) was associated with a significant reduction in CNS extracellular K^+^ concentration compared with w1118 flies (n=23, median 7.6 mM, IQR 6.1-8.9 mM) (*P* < 0.01, Mann-Whitney test) (Figure 8B).

**Figure 8.**
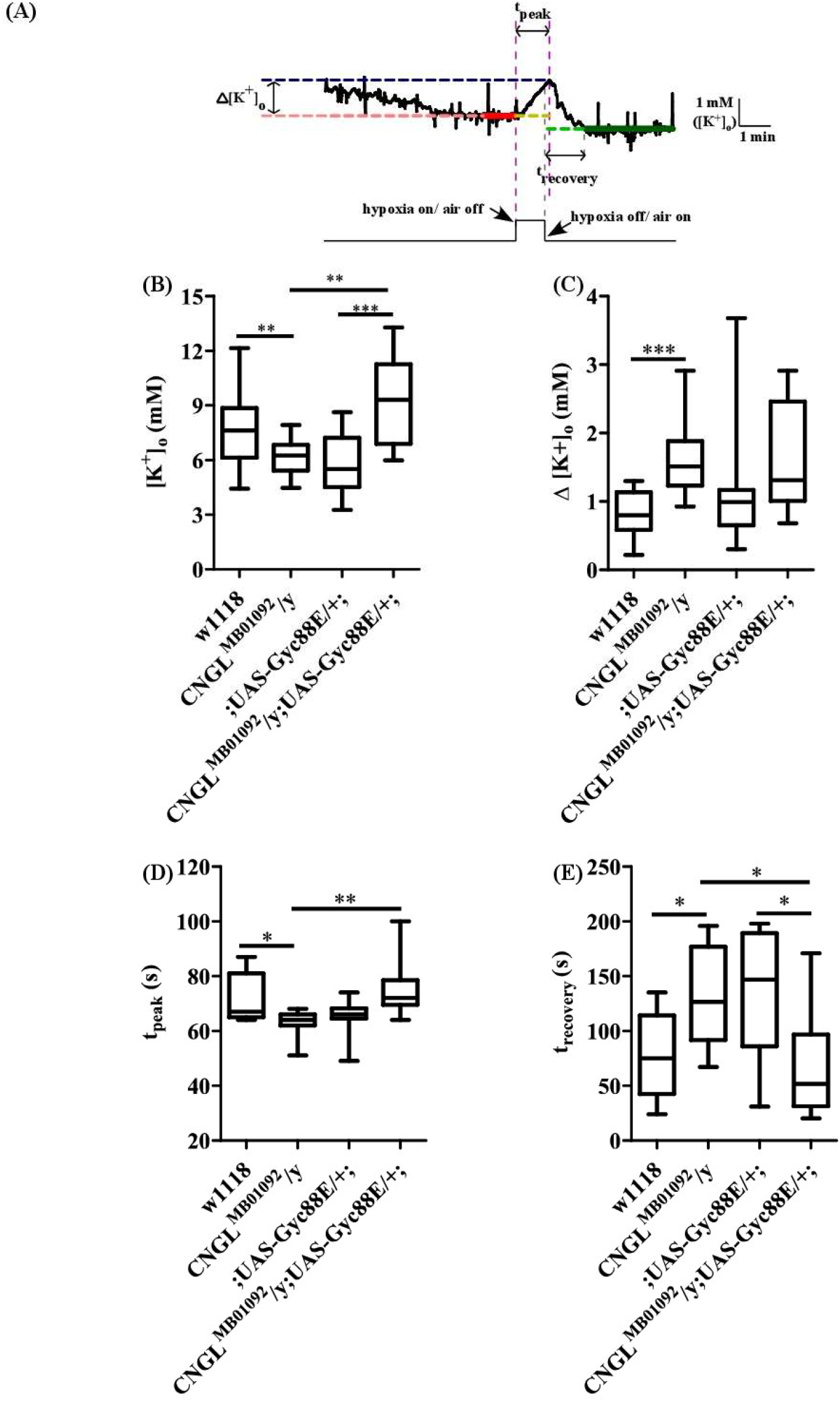
Reduced basal levels of CNS extracellular K^+^, and increased ∆ [K^+^]o and trecovery, and reduced tpeak in response to hypoxia in CNGL^MB01092^/y flies. The overexpression of Gyc88E eliminated the reduced basal levels of of CNS extracellular K^+^ and partially compensated for the CNS alterations in response to hypoxia in flies with down-regulation of *CNGL*. (A) The response variables in the [K^+^]o trace. The red and green lines represent the baselines before and after hypoxia, respectively. The blue line represents the peak concentration. (B) Basal levels of CNS extracellular K^+^ of w1118 (n=23), CNGL^MB01092^/y (n=19),;UAS-Gyc88E/+; (n=14), and CNGL^MB01092^/y; UAS-Gyc88E/+; (n=12). (C-E) The parameters of ∆[K^+^]o, tpeak and trecovery in w1118 flies (n=9), CNGL^MB01092^/y flies (n=10),;UAS-Gyc88E/+; (n=14), and CNGL^MB01092^/y;UAS-Gyc88E/+; (n=10). Asterisks (*, ** or ***) indicate *P* < 0.05, 0.01 or 0.001, respectively, by Mann-Whitney test or Kruskal-Wallis test with Dunn’s multiple comparison as indicated in the Results.

### 3.9 Overexpression of Gyc88E compensated for the reduced basal levels of CNS extracellular K^+^ in *CNGL* mutant

cGMP upregulation by the overexpression of Gyc88E eliminated the reduced locomotion in *CNGL* mutant flies. Thus we examined whether it could also suppress the reduced basal levels of CNS extracellular K^+^ in CNGL^MB01092^/y flies.

Gyc88E overexpression led to a marked baseline increase in CNS extracellular K^+^ recording (*P* < 0.01 or 0.001, Kruskal-Wallis test with Dunn’s multiple comparison) (Figure 8B). [K^+^]_o_ in flies CNGL^MB01092^/y; UAS-Gyc88E/+; was 9.3 mM (n=12, IQR 6.9-11.3 mM), however, the two control flies, CNGL^MB01092^/y and;UAS-Gyc88E/+;, had K^+^ baseline at 6.2 mM (n=19, IQR 5.4-6.8 mM) and 5.5 mM (n=14, IQR 4.5-7.2 mM), respectively. Additionally, the basal level of CNS extracellular K^+^ in CNGL^MB01092^/y; UAS-Gyc88E/+; flies was also significantly higher than that of w1118 flies (*P* < 0.05, Mann-Whitney test), suggesting that overexpression of Gyc88E eliminated the reduced basal levels of CNS extracellular K^+^ and even increased it to a higher level in *CNGL* mutant.

### 3.10 Increased ∆ [K^+^]o and t_recovery_, and reduced t_peak_ in CNGL^MB01092^/y flies in response to hypoxia

Hypoxia caused a disturbance in ion homeostasis in the brain. [K+]_o_ increased during hypoxia and returned to baseline when normoxia was restored (Figure 8A). Hypoxia caused a larger change in [K+]_o_ (∆[K+]_o_) in CNGL_MB01092_/y flies (n=10, median 1.5 mM, IQR 1.2-1.9 mM) compared with w1118 flies (n=9, median 0.8 mM, IQR 0.6-1.1 mM) (*P* < 0.001, Mann-Whitney test) (Figure 8C). In addition, the mutant flies took significantly shorter times to reach the peak. The t_peak_ of CNGL^MB01092^/y flies was 64.0 s (n=10, IQR 62.0-66.0 s) while that of control flies was 67.0 s (n=9, IQR 65.0-81.0 s) (*P* < 0.05, Mann-Whitney test) (Figure 8D). The other parameter we measured was t_recovery_. CNGL^MB01092^/y flies (n=10, median 126.5 s, IQR 91.8-176.8 s) took longer to regain their baseline compared with w1118 flies (n=9, median 75.0 s, IQR 42.5-114.0 s) (*P* < 0.05, Mann-Whitney test) (Figure 8E).

Therefore, the effect of hypoxia on K^+^ homeostasis was more severe in *CNGL* mutant fly line CNGL^MB01092^/y by showing an increased ∆[K^+^]_o_ and t_recovery_, and a reduced t_peak_ compared with w1118 control flies.

### 3.11 Overexpression of Gyc88E significantly reduced t_recovery_ in *CNGL* mutant in response to hypoxia

The current study has shown that the overexpressing Gyc88E compensated for the reduced locomotion in CNGL mutant after hypoxia. We then tested whether it could also eliminate the disruption to ion homeostasis in response to hypoxia in CNGL mutant flies.

∆[K^+^]_o_ was similar between flies CNGL^MB01092^/y;UAS-Gyc88E/+; (n=10, median 1.3 mM, IQR 1.0-2.5 mM) and;UAS-Gyc88E/+; (n=14, median 1.0 mM, IQR 0.7-1.2 mM) (*P* > 0.05, Kruskal-Wallis test with Dunn’s multiple comparison) (Figure 8C). Although hypoxia caused a lower ∆[K^+^]_o_ when Gyc88E was overexpressed in *CNGL* mutant compared with CNGL^MB01092^/y flies (n=10, median 1.5 mM, IQR 1.2-1.9 mM), no statistically significant difference was observed (*P* > 0.05, Kruskal-Wallis test with Dunn’s multiple comparison).

The t_peak_ of CNGL^MB01092^/y;UAS-Gyc88E/+; was 72.0 s (n=10, IQR 69.5-78.5 s). Although it was similar to that of;UAS-Gyc88E/+; flies (n=14, median 66.0 s, IQR 64.5-68.3 s), it was higher than that of *CNGL* mutant flies (n=10, median 64.0 s, IQR 62.0-66.0 s) (*P* > 0.05 or < 0.01, respectively, Kruskal-Wallis test with Dunn’s multiple comparison) (Figure 8D). In addition, flies CNGL^MB01092^/y;UAS-Gyc88E/+; (n=10, median 51.5 s, IQR 31.3-96.8 s) required less time to recover from hypoxia compared with two control fly lines (CNGL^MB01092^/y, n=10, median 126.5 s, IQR 91.8-176.8 s; and;UAS-Gyc88E/+;, n=14, median 147.0 s, IQR 86.0-189.3 s) (*P* < 0.05 for both comparisons, Kruskal-Wallis test with Dunn’s multiple comparison) (Figure 8E).

Hence, the results suggest that in response to hypoxia, the upregulation of cGMP by the overexpression of Gyc88E significantly reduced t_recovery_ in *CNGL* mutant.

## 4 Discussion

In control w1118 flies, hypoxia immediately reduced locomotor performance measured by path length per minute, however there was complete recovery upon the return of air (~40 mins to recover after 15 mins of hypoxia). CNGL^MB01092^/y mutant flies had low levels of *CNGL* transcripts and reduced locomotion in normoxia with poor recovery from hypoxia. Reduced locomotor performance in the mutant was a result of more time spent motionless in between episodes of walking and a lower walking speed during episodes. CNGL^MB01092^/y flies also had an immediate increase in path length per minute at the beginning of hypoxia, followed by a gradual increase during the hypoxia period. *CNGL* knockdown in neurons replicated the properties of the CNGL^MB01092^/y mutant. However, *CNGL* knockdown in glial cells impaired locomotion to such an extent that it was not possible to characterize effects of hypoxia and return to air. The CNGL^MB01092^/y mutant had reduced basal levels of extracellular K^+^ concentrations in the brain compared with wild-type w1118 flies. Downregulation of *CNGL* in the mutant also affected the dynamics of [K^+^]_o_ in response to brief hypoxia by reducing the time to reach a higher peak [K^+^]_o_ and increasing the time to recover to baseline. Finally, we showed that crossing the mutant with UAS-Gyc88E or the Pde1c^KG05572^ mutant, to upregulate cGMP, compensated for *CNGL* downregulation and enabled locomotor recovery from hypoxia in the mutant line. Gyc88E also partially compensated for the ion homeostasis differences between w1118 controls and *CNGL* mutants.

The interpretation of our results is limited in two major ways. First, although we show that *CNGL* transcript level is lower in the CNGL^MB01092^/y mutants, we do not know the effect of this on channel abundance, kinetics or ion currents. Our conclusions therefore relate to the clear and undeniable effects of this particular genetic mutation on responses to hypoxia rather than to a defined manipulation of channel function. We speculate that CNGL channel activity is reduced in the mutant but this requires confirmation in future experiments. Second, for technical reasons we did not outcross the fly lines a sufficient number of times to be certain that our results were not affected by different genetic backgrounds. Thus, it remains a formal possibility that differing genetic backgrounds influenced the results. However, the effects of a mutation that reduced levels of *CNGL* transcript were recapitulated by RNAi targeting *CNGL* in neurons (and possibly glia) and were compensated by two separate manipulations (Gyc88E and Pde1c) that would be expected to increase cGMP levels. Thus, we argue that the most parsimonious explanation that fits all of our data and previous published data on CNG channels, responses to hypoxia in larvae and the role of a *white*/cGMP/PKG signaling pathway in responses of adult flies to anoxia (11, 62, 66, 73), is that our genetic manipulations had their intended effects.

We found that CNGL^MB01092^/y had reduced locomotion under normoxia, suggesting that *CNGL* is necessary for normal motor activity in adult flies. This notion is supported by the structural similarity of CNGL with HCN channels (74), which are intimately involved in the central generation of motor patterns in many taxa (50). Indeed, it is intriguing that many of the effects of *CNGL* mutation on fly walking are similar to the effects of blocking HCN channels on swimming of larval *Xenopus*, including shorter episodes of slower locomotion. In larval *Xenopus*, HCN currents compensate for the hyperpolarizing activity of the Na^+^/K^+^-ATPase linking the locomotor effects to ion homeostasis in the central circuitry (50). The circuitry for walking in flies is unknown but it is tempting to speculate that CNGL channels may have modulatory influences on interneuronal activity and interactions similar to HCN channels in tadpoles.

Compared with w1118 control flies, down-regulation of *CNGL* also led to an 18.4% reduction in extracellular K^+^ concentration. A possible explanation for the reduced locomotion in *CNGL* mutant flies is that the reduced extracellular K^+^ concentration would increase the K^+^ equilibrium potential, thus hyperpolarizing neurons and reducing excitability. Moreover, K^+^ conductance and currents clearly affect motor activity in animals. For example, protein tyrosine phosphatase alpha (PTPα) stimulates voltage-gated potassium Kv1.2 activity (63), and PTPα knockout mice show decreased exploratory locomotor activity (59). In addition, the application of 5-Hydroxytryptamine (5-HT), which reduces potassium conductance in rat (49), decreases locomotor activity (51). We did not monitor other ions and changes of Na^+^ and Ca^2+^ concentrations might also contribute to the altered locomotion in *CNGL* mutant flies. In *Drosophila* larvae, mutations in Pickpocket1 (PPK1), which belongs to the sodium channel family, enhance locomotion by increasing crawling speed and decreasing stops and turns relative to wild-type (1). Voltage-dependent Ca^2+^ channel (Ca_v_2.2) knockout mice show more activity than controls (Beuckmann et al., 2003).

The hypoxia escape response in *Drosophila* larvae requires functional CNGA channels but not CNGL channels (66). This is consistent with the observation that CNGA channels need to be activated for the mobilization of downstream target Ca^2+^, however, CNGL channels are not involved (13). cGMP modulates the speed of anoxic recovery in adult flies (73) and mediates the escape response to hypoxia in *Drosophila* larvae (66). One of the cGMP downstream targets, PKG, is involved in behavioral tolerance to anoxia and hypoxia (11, 62). The current study shows that the CNGL channel, one member of the CNG ion channel family, which represents downstream targets of cGMP, also regulated the hypoxia response in adult flies. However, whether the hypoxia response in adults also requires the other members of CNG channels such as CNGA, was not investigated in our research. The mechanism of sensory neuron activation in response to hypoxia is proposed in *Drosophila* larvae (46). In larvae, hypoxia activates two heterodimeric atypical sGCs (Gyc-88E/Gyc-89Da and Gyc-88E/Gyc-89Db), which act as O_2_ sensors. The sGCs then promote the production of cGMP, which activates CNG channels. *Drosophila* larvae withdraw from food in response to hypoxia (66) or remain moving in anoxia for almost 40 min (5), whereas adults show complete paralysis within 30 s under anoxia (5). The different behavioral responses to hypoxia in larvae and adults suggest that different mechanisms of responding to hypoxia are involved and therefore, whether O_2_ sensors are required in hypoxia response of adults is unclear. To examine the involvement of O_2_ sensors in adult flies, the hypoxia responses in the atypical sGCs mutant flies should be compared with the controls.

Under normoxia, one of the factors leading to the reduced path length per minute in CNGL^MB01092^/y flies was the increased times spent motionless between episodes of walking. In larval zebrafish, fictive swimming with an episodic pattern is evoked by the application of N-methyl-d-aspartate (NMDA), which is an activator of the larval zebrafish swimming central pattern generator (42, 67). When *C.elegans* swim, episodic locomotion is also observed with spontaneous alternation between active swimming and inactivity, and this transition is promoted by acetylcholine (ACh) signaling (19). So far, no direct evidence has been found for the involvement of the CNGL channel in the release of NMDA or ACh, however, it is reported that the CNG channels are able to modulate the release of glutamate (4, 55, 75). Therefore, under normoxia, the increased duration of quiescence with fewer episodes of walking may have been due to a disruption of neuromodulation resulting from the down-regulation of *CNGL*.

The behavioral response to hypoxia in *Drosophila* larvae depends on the nitric oxide (NO)/cyclic GMP (cGMP) signaling pathway (68). In swimming *Xenopus laevis* larvae the application of S-nitroso-N-acetylpenicillamine (SNAP) to increase NO levels, results in a reduced episode duration (the time for which animal swims in response to a brief sensory stimulus) and an increased cycle period (interval between the onset of a burst in one cycle and the onset of the burst in the next cycle) (43). On the other hand, treatment with nitric oxide synthase (NOS) inhibitors, such as L-NAME and L-NNA, to reduce NO levels increases the episode duration and decreases the cycle period. The effect of SNAP on the swimming motor pattern is similar to that of sodium azide (NaN_3_) when it is applied to *Xenopus laevis* larvae (52), which is widely used to induce chemical hypoxia in neurons (32, 56, 65) and is reported to generate NO (60, 61). In adult flies, it is still unclear why an acute increase of locomotion was generated at the beginning of hypoxia, especially in the flies with *CNGL* down-regulation or pan-neuronal knockdown. However, it demonstrates that the locomotor system is capable of operating faster. Reduced locomotion followed by the acute increase value in the fly lines noted above as well as the decreased activity in all other tested fly lines during hypoxia might be due to increased levels of NO, reducing episode duration and thus decreasing the path length per minute.

Astrocytes, a type of glial cell, are critical for the behaviors in *Drosophila* larvae by regulating the Ca^2+^ level and the downstream dopaminergic neurons (41). Octopamine/tyramine receptor (Oct-TyrR) and the transient receptor potential (TRP) channel Water witch (Wtrw) are essential in this astrocytic Ca^2+^ signaling pathway and are required for normal behavior in larvae. Under normoxia, flies with *CNGL* knocked down in glia showed remarkably reduced locomotor performance compared both with control flies and even with flies with *CNGL* down-regulation or knockdown in neurons. It is possible that the CNGL channel is also essential in the glial Ca^2+^ signaling pathway and critical for the normal neural operation and locomotion in adult flies. CNGL^MB01092^/y flies showed reduced recovery after hypoxia. This was comparable with *CNGL* knockdown in neurons, however, the comparison with *CNGL* knockdown in glia was not possible because of impaired performance in these flies. The *CNGL* gene is highly expressed in neurons (44), and it is likely that the CNGL channels in glial cells have different functions from the CNGL channels in neurons, leading to the different sensitivities to hypoxia or varied effects of O_2_ level changes on locomotor activities. Nevertheless, compared with Gal4 control flies, the reduced recovery of flies with *CNGL* knockdown in neurons suggests that CNGL channels expressed in neurons regulated hypoxic recovery. To determine the contributions of specific neuronal or glial cell types to the hypoxia response, specific driver lines for neurons or glia of the fly CNS could be used to cross with UAS-CNGL-RNAi flies.

We also found that the overexpression of Gyc88E or the mutation of *Pde1c* compensated for the reduced recovery from hypoxia in *CNGL* mutant flies. We did not measure cGMP levels when Gyc88E was overexpressed, however, the overexpression of Gyc88E eliminates the *white* (*w*) gene-RNAi-induced delay of locomotor recovery and the effect is similar to several PDE mutants, especially cGMP-specific PDE mutants (73), suggesting that the Gyc88E overexpression is likely to have increased cGMP levels. Genes for cGMP-specific PDEs, which also regulate intracellular levels of cGMP by hydrolyzing cGMP, have been identified in *Drosophila* (12, 47). Therefore, *PDE* mutation is another approach to investigate the impact of cGMP upregulation (73). One of the *PDE* mutants, *Pde1c*^KG05572^, could compensate for the *w*-RNAi-induced delay of locomotor recovery from anoxia. It should be noted that;UAS-Gyc88E/+; recovered poorly from hypoxia. The reason for this inconsistency between these two approaches might be that *Pde1c* mutation was not a UAS-line and therefore, it lost tissue specificity and could not function as specifically as the UAS-Gyc88E fly line, in which Gyc88E was overexpressed only in *CNGL*-positive cells. The reason for this is unclear, however, compared with *CNGL* mutant flies CNGL^MB01092^/y, the remarkable increased recovery of the flies with Gyc88E overexpression or *Pde1c* mutation in *CNGL* mutants suggests that cGMP upregulation eliminated the impaired recovery induced by the down-regulation of *CNGL* in response to hypoxia.

Another interesting observation is that post-hypoxic depression of locomotion was observed in the mutant flies as well as w1118 flies. This is very noticeable in the CNGL^MB01092^/y; *Pde1c*^KG05572^/+; flies (Figure 6C,D) and appears similar to the post-hypoxic depression of respiratory rhythm generation in male mice (18). At the onset of reoxygenation, a large peak of CO_2_ output, which is an index of mitochondrial activity, can be observed in male flies, and this peak reduces with increased anoxia duration (36). In addition, the increased CO_2_ emission corresponds to a lack of activity at the beginning of reperfusion with O_2_ in flies (36). Mitochondria produce reactive oxygen species (ROS) including superoxide (17, 39). Therefore, the initial trigger for the post-hypoxic depression of locomotion is likely due to the resumed function of mitochondria and the increased level of superoxide. Moreover, mitochondria also produce nitric oxide (NO) (16), which stimulates sGCs and increases cGMP production. The full recovery from hypoxia depends on the clearance of superoxide and metabolites such as adenosine, due to their effects on locomotor neural circuits and muscles (20, 69).

The electrophysiological results in the current study showed that in response to hypoxia, the down-regulation of *CNGL* led to changes in the dynamics of [K^+^]_o_ in the brain suggesting that the down-regulation of *CNGL* altered extracellular ion homeostasis. Anoxia causes a surge of [K^+^]_o_ in fly brain (3). The increased ∆[K^+^]_o_ in response to hypoxia indicated that compared with w1118 control flies, ion homeostasis in *CNGL* mutant, especially with respect to [K^+^]_o_, was more easily disrupted and these flies might have difficulty in maintaining [K^+^]_o_ homeostasis in response to hypoxia. *CNGL* mutant flies required shorter time to reach the [K^+^]_o_ peak, which also indicates that they had a more severe disruptive effect on [K^+^]_o_ homeostasis. Taking t_peak_ as another measure of ion homeostasis and an indirect measure of hypoxia tolerance, our result provided further evidence that the down-regulation of *CNGL* disturbed ion homeostasis under hypoxia. It is also worth noting that most of the flies reached [K^+^]_o_ peak after hypoxia was turned off, resulting in t_peak_ larger than the hypoxia duration which was 60 s. The reason might be that flies required a short while to respond to the increased O_2_ level, which was consistent with previous studies (3, 54).

In summary, our results support the conclusion that the CNGL channel is involved in ion homeostasis in the brain and the recovery from hypoxia in *Drosophila* adults. With respect to central pattern generation, the role of cGMP and CNGL channels in episodic motor activity and post-hypoxic inhibition are future research areas that are likely to be particularly fruitful.

## Acknowledgement

Funded by a Discovery Grant from the Natural Sciences and Engineering Research Council of Canada (#40930-2009).

